# Cell Communication Network factor 4 promotes tumor-induced immunosuppression in breast cancer

**DOI:** 10.64898/2026.01.07.698253

**Authors:** David J. Klinke, Atefeh Razazan, Anika C. Pirkey, Wentao Deng

## Abstract

Cell Communication Network factor 4 (CCN4/WISP1) is a matricellular protein secreted by cancer cells that is upregulated in essentially all invasive breast cancers and promotes immunosuppression in melanoma. Recent work suggests that limited anti-tumor immunity also associates with poor patient outcomes in patients with breast cancer. Motivated by increased CCN4 correlating with dampened anti-tumor immunity in primary breast cancer, we test for a direct causal link by knocking out CCN4 (CCN4 KO) in the Py230 and Py8119 mouse breast cancer models. Tumor growth is reduced when CCN4 KO breast cancer cells are implanted in immunocompetent but not in immunodeficient mice. Correspondingly in size-matched tumors, CD4+ and CD8+ T cells are significantly increased in CCN4 KO tumors while the myeloid compartment is shifted from polymorphonuclear- to monocytic-myeloid-derived suppressor cells (MDSC). This shift in the MDSC compartment is also associated with a significant reduction in splenomegaly. Among mechanisms linked to local immunosuppression, CCN4 knockout has a similar impact on the secretome of both breast cancer and melanoma cell lines. Overall, our results suggest that CCN4 promotes tumor-induced immunosuppression in the context of breast cancer and is a potential target for therapeutic combinations with immunotherapies.

## Introduction

Breast cancer remains a leading cause of cancer mortality in women worldwide, with considerable heterogeneity in patient outcomes that cannot be easily mapped onto traditional histopathological classifications. While breast cancer has long been considered a poorly immunogenic malignancy, observations that tumor-infiltrating lymphocytes are enriched in the tumor microenvironment (1) presage the clinical successes of immune checkpoint blockade in patients with advanced triple-negative breast cancer (2). Still, the majority of breast cancer patients do not respond to current immunotherapies, highlighting a significant unmet medical need. One way to expand the clinical impact of immunotherapies is to identify the biochemical signals that malignant cells use to evade host immunosurveillance and reshape the tumor microenvironment to suppress anti-tumor immunity.

Cell Communication Network factor 4 (CCN4), also known as WNT1-inducible signaling pathway protein 1 (WISP1), is a secreted matricellular protein that is essentially upregulated in all patients with invasive breast cancer (3). Matricellular proteins function not as structural components of the extracellular matrix but rather as modulators of cell-matrix interactions and cellular responses to environmental cues. In the context of melanoma, CCN4 promotes both metastasis via the induction of the epithelial-mesenchymal transition (EMT) and tumor-induced immunosuppression (4–6). The EMT represents a coordinated developmental program whereby epithelial cells acquire mesenchymal characteristics, including enhanced motility, invasiveness, and resistant to apoptosis. In terms of gene expression patterns linked to the EMT, a similar shift in gene expression occurs in human melanoma and breast cancer (7). Interestingly, CCN4 was one of only three secreted proteins that were associated with a mesenchymal-like state in both cancer types, suggesting conserved functional roles that transcend tissue-specific contexts.

Prior bioinformatic analysis of primary breast cancer samples revealed that increased CCN4 expression correlates with dampened anti-tumor immunity, as evidenced by reduced signatures of type 1 cell-mediated cytotoxic immunity (3). More recent Bayesian network analysis predicted that CCN4 has a similar impact on the prevalence and functional orientation of immune cells within the tumor microenvironment in the context of melanoma and breast cancer (8). While Bayesian networks offer causal interpretations of human data, they are still predictions that require experimental validation. Given that CCN4 promotes tumor-induced immunosuppression in melanoma through alterations in the secretome (6), and given the shared gene expression patterns between melanoma and breast cancer (7), we hypothesized that CCN4 directly suppresses anti-tumor immunity in breast cancer.

The objective of this study is to test for a direct causal link by knocking out CCN4 (CCN4 KO) in the Py230 and Py8119 cell lines, two transplantable immunocompetent mouse models of breast cancer. These cell lines, derived from MMTV-PyMT transgenic mice, recapitulate key features of human breast cancer and permit evaluation of tumor-immune interactions in syngeneic, immunocompetent hosts (9). By comparing tumor growth dynamics in immunocompetent versus immunodeficient mice, characterizing tumor-infiltrating lymphocyte profiles in size-matched tumors, quantifying systemic immune perturbations, and profiling the cancer cell secretome, we demonstrate that CCN4 actively promotes immunosuppression in breast cancer through mechanisms that parallel those observed in melanoma.

## Results and Discussion

### CCN4 knockout reduces tumor growth in an immune-dependent manner

To test whether CCN4 directly promotes immunosuppression in breast cancer, we generated CCN4 knockout (CCN4 KO) variants of two transplantable mouse breast cancer cell lines, Py230 and Py8119, using CRISPR/Cas9 double nickase systems. The Py230 and Py8119 parental cells secreted 1185 *±* 72 pg/ml and 243 *±* 188 pg/ml, respectively, of CCN4 in 2D culture media. Of note, CCN4 production by Py230 cells was similar to that observed by WT B16F0 cells (1015 *±* 50 pg/ml). Two independent knockout clones were isolated for each parental line and validated by ELISA. These CCN4 KO variants produced undetectable levels of CCN4 under similar culture conditions (Supplemental Figure S1).

Following subcutaneous (s.c.) implantation, we compared the tumor growth trajectories between WT and CCN4 KO variants in immunocompetent C57BL/6 mice to their growth in severely immunocompromised NSG mice. In NSG mice that lack functional T, B, and NK cells, CCN4 knockout had minimal impact on tumor development for both Py8119 and Py230 cell lines (Figure 1A, C). In contrast, CCN4 KO significantly delayed tumor onset in immunocompetent C57BL/6 mice and improved survival for both models (Figure 1B, D). For Py8119 variants, the difference was highly significant (p-value = 3e-4), while the Py230 variants showed an even more pronounced effect (p-value *<* 1e-7). In fact, all 10 C57BL/6 mice challenged with Py230 CCN4 KO clone 2 failed to develop tumors. These results demonstrate that CCN4’s tumor-promoting effects require an intact immune systems and support a direct role for CCN4 in mediating tumor-induced immunosuppression rather than purely cancer cell-intrinsic effects on proliferation or survival. While we note that subcutaneous injection does not fully recapitulate the mammary gland microenvironment, the fundamental question we addressed whether CCN4 promotes immunosuppression - is unlikely to depend on the anatomical site of tumor growth. Preliminary studies using orthotopic injection into the mammary gland (5/5 C57BL/6 mice developed WT Py230 tumors while only 1/5 mice developed Py230 CCN4 KO1 tumors and 0/5 mice developed Py230 CCN4 KO2 tumors after 33 days) were consistent with the s.c. results presented here but quantifying the tumor growth trajectories were more variable than we anticipated. Given the focus on size-matched experiments, we relied on subcutaneous implantation for subsequent analysis.

**Fig. 1.**
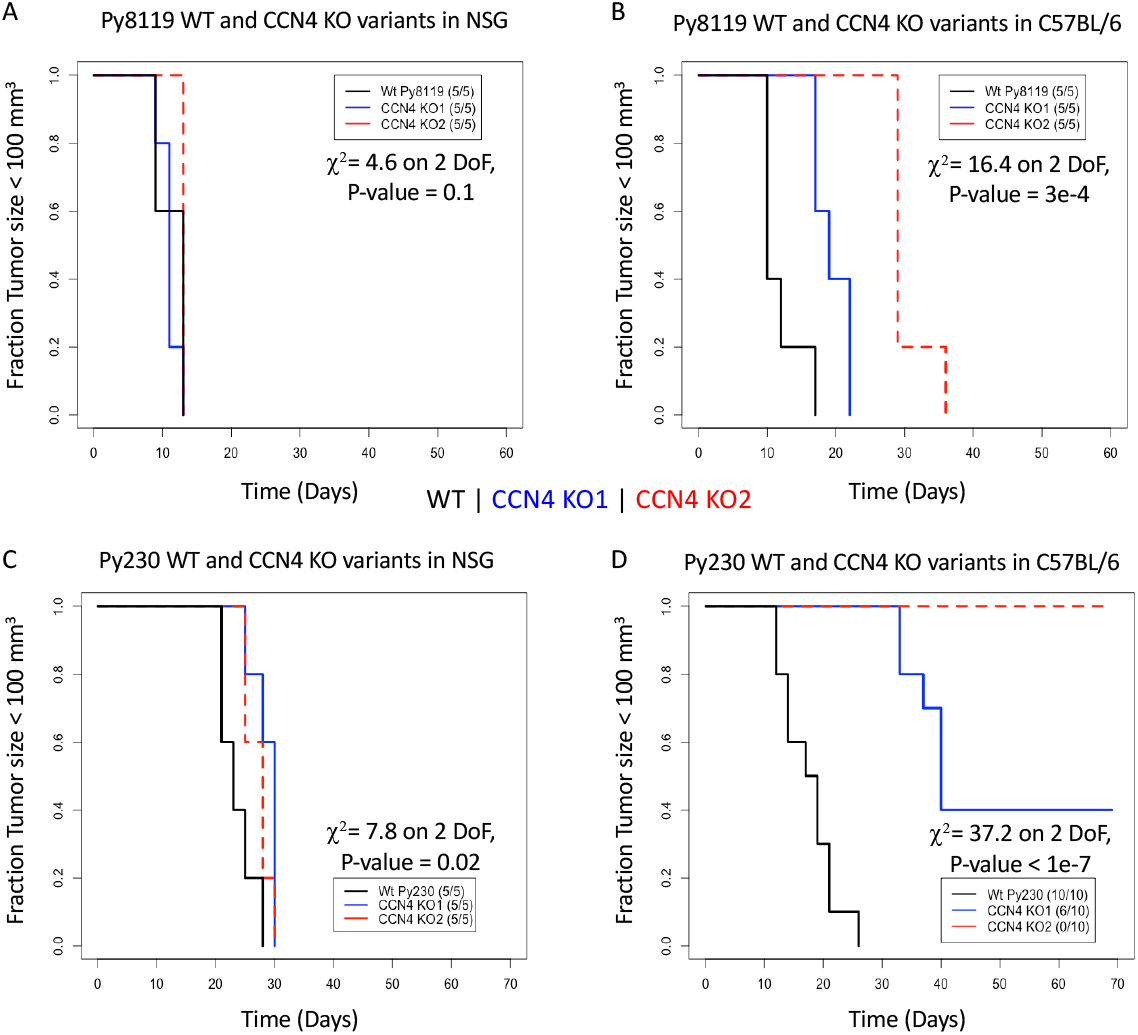
CCN4 KO has minimal impact on immunocompromised NSG mice but improves survival of immunocompetent C57BL/6 mice challenged with Py230 and Py8119 variants. Kaplan-Meier time-to-event survival curves for immunocompetent C57BL/6 mice (B and D) and severely immunocompromised NSG mice (A and C) challenged subcutaneously with wild-type (black curves) and two CCN4 KO variants (KO1 in blue and KO2 in dotted red curves) derived from Py8119 (A and B) and Py230 (C and D) cell lines. The tumor size exceeding 100 mm^3^ was considered the triggering event. The number of animals that experienced a triggering event among the total number of animals challenged with tumor cells is indicated in the legend. Statistical difference among the three curves was assessed using a Peto & Peto Modification the Gehan-Wilcoxon test with the resulting Chi-squared, degrees of freedom and resulting P-values are indicated. As this was an exploratory study, the p-values indicate the extent to which these three Kaplan-Meier survival curves are different.

### CCN4 knockout reduces systemic immunosuppression

Splenomegaly, or enlargement of the spleen, often accompanies tumor-induced systemic immunosuppression and correlates with the expansion of immunosuppressive myeloid populations (10). In a time-matched experimental design, mice bearing WT Py230 tumors exhibited significantly greater splenomegaly than those challenged with CCN4 KO tumors (p-value *<* 0.001, Figure 2A). As implied by these results and similarly described by others (11), tumor-burden alone can be an independent predictor of splenomegaly in tumor cell transplantable models. To rule out that possibility, we used a size-matched experimental design with Py8119 tumors. In those experiments, WT tumor-bearing mice showed significantly greater splenomegaly than those with CCN4 KO tumors (p-value = 0.011, Figure 2B) despite similar tumor burdens. The reduction in splenomegaly upon CCN4 knockout suggests that CCN4 contributes to systemic immune perturbations that extend beyond the local tumor microenvironment.

**Fig. 2.**
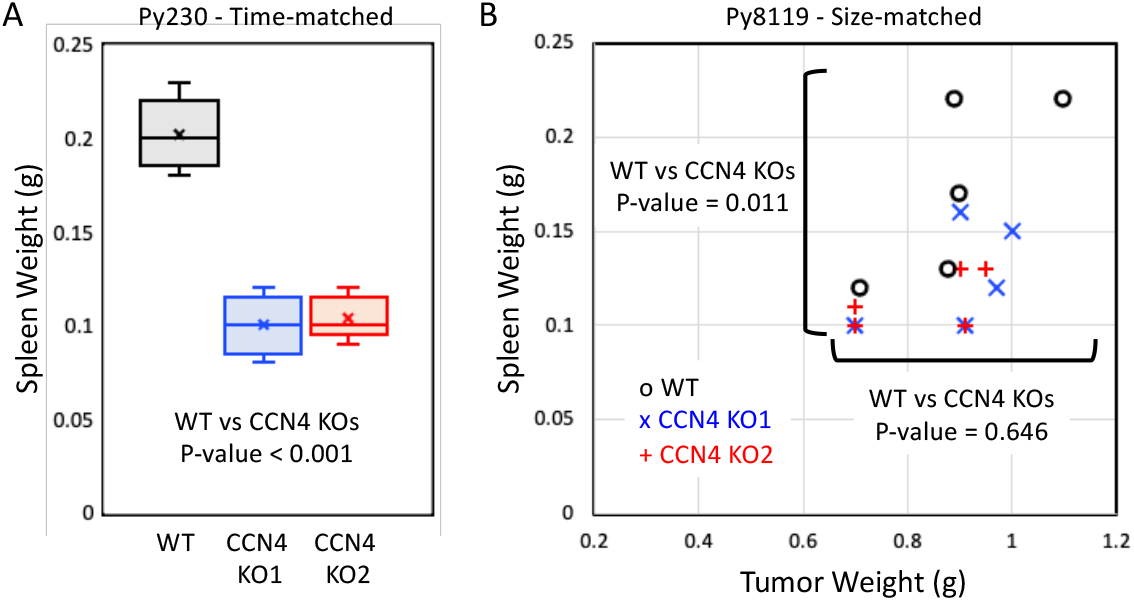
Splenomegaly occurs in mice challenged with WT Py230 and Py8119 cells compared to CCN4 KO variants. (A) Box-and-whisker plots for spleen weights obtained from C57BL/6 mice challenged with WT Py230 cells and CCN4 KO variants In a time-matched experimental design with the end point at 69 days (n = 5 for each group). (B) Spleen weights and corresponding excised tumor weights obtained from C57BL/6 mice challenged with WT Py8119 cells and CCN4 KO variants in a size-matched experimental design (n = 5 for each group). As there was no statistical difference between CCN4 KO variants, results were combined for analysis. A homoscedastic two-sided Student’s t-test was used to assess statistical significance.

### CCN4 knockout enhances T cell infiltration and alters myeloid compartment

To characterize how CCN4 shapes the tumor immune microenvironment, we performed flow cytometric analysis of tumor-infiltrating lymphocytes (TILs) in size-matched tumors. Using full-spectrum flow cytometry and self-organizing maps to visualize the immune landscape in tumors approximately 300 mm^3^ in size, we focused on the nodes that exhibited high concordance with annotations obtained using manual gating (Figure 3). We found that CCN4 KO1 Py230 tumors exhibited increased infiltration of both CD4+ and CD8+ T cells (Live CD45+ CD3e+ CD4+ CD8aFoxp3-CD25- and Live CD45+ CD3e+ CD4-CD8a+ events) compared to WT tumors (Figure 3). Correspondingly, the overall presence of myeloid-derived suppressor cells (MDSCs), which actively suppress anti-tumor T cells response, were reduced in CCN4 KO tumors. In addition, there was a compositional shift in the myeloid compartment from the prevalence of polymorphonuclear MDSCs (PMN-MDSC : Live CD45+ CD3e- CD11b+ CD11c- Ly-6G+ Ly- 6C^*int*^ MHCII- F4/80-) and tumor-associated macrophages (Live CD45+ CD3e- CD11b- CD11c+ Ly-6C-) to monocytic MDSCs (Mo-MDSC: Live CD45+ CD3e- CD11b+ CD11c- Ly-6C+ Ly-6G- MHCII-) upon CCN4 KO.

**Fig. 3.**
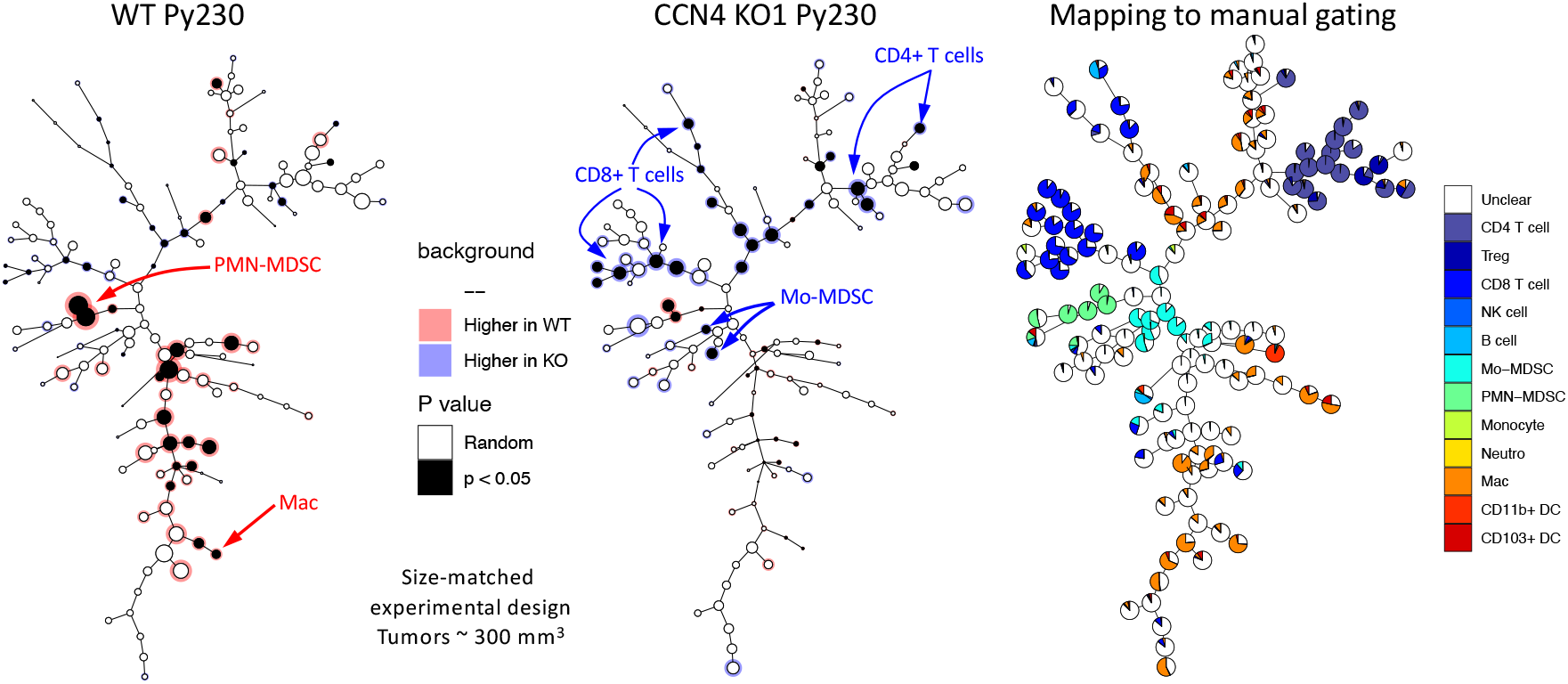
T cells are elevated and MDSCs are reduced in CCN4 KO1 compared to WT Py230 tumors. Following enzymatic dissociation of tumor tissues, the immune cell compartment was quantified by full spectrum flow cytometry. A self-organizing map shows the distribution of negative Live/Dead violet and positive CD45 cells in size-matched WT (left) and CCN4 KO1 (center) tumors (n = 5 for both WT and CCN4 KO1 Py230). Right panel shows the percentage of events associated with each cluster that relate to cells identified by manual gating.

In the Py8119 model, we observed similar patterns using two different antibody panels that focused on T, B, and NK cells and myeloid cell subsets instead of full-spectrum flow cytometry. In this size-matched experiment, tumor sizes for the corresponding cohorts are reported in Figure 2B. While total CD4+ T cell infiltration was not significantly different (p-value = 0.133, Figure 4B), the CD25+ fraction of CD4+ T cells was significantly elevated in CCN4 KO tumors (p-value *<* 0.001, Figure 4C). CD25 expression on CD4+ T cells can mark both activated effector T cells and regulatory T cells (Tregs), suggesting altered T cell activation states upon CCN4 loss. While these two T cell populations have opposing effects on anti-tumor immunity, we note that the full-spectrum flow cytometry analysis of Py230 tumors included FoxP3 staining and suggested that the Treg population was not substantially altered (Figure 3), implying that the increased CD25+ CD4+ T cells likely represent activated effector cells.

**Fig. 4.**
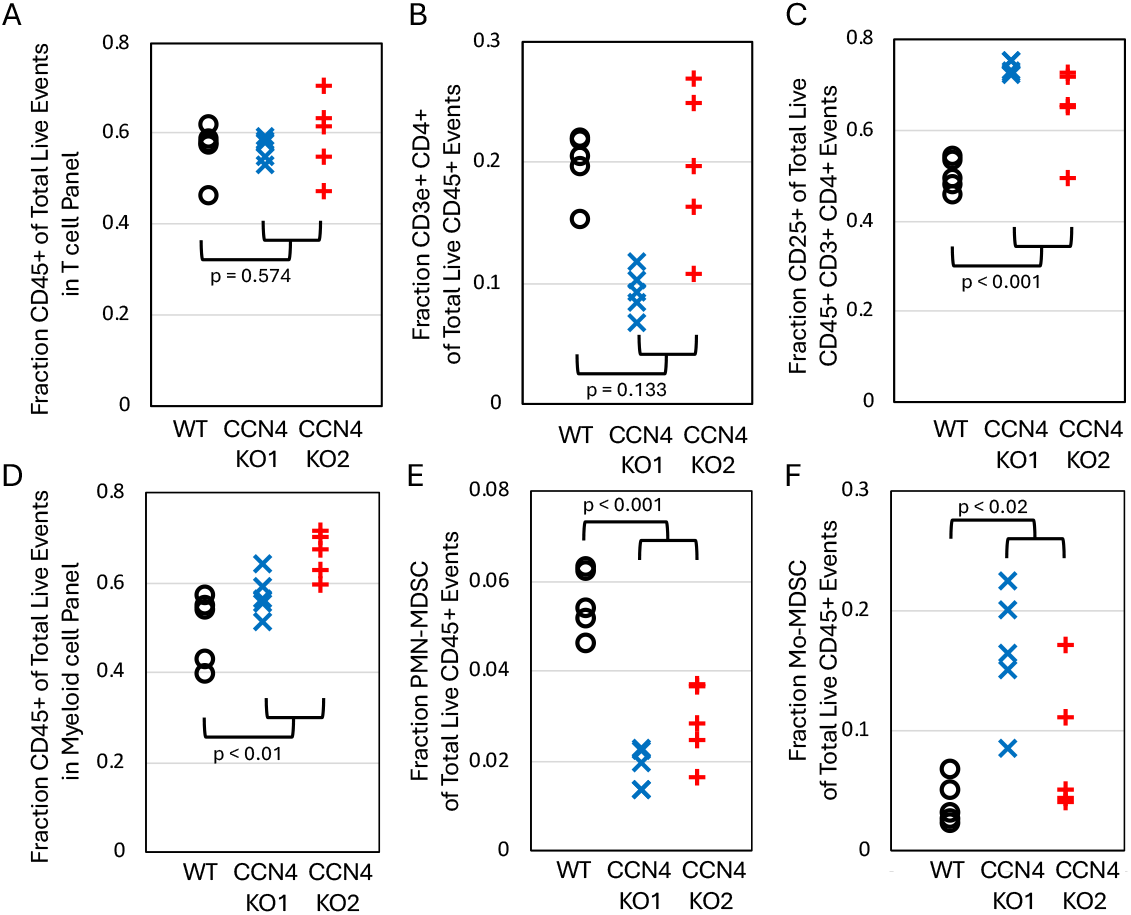
CD25+ CD4+ T cells are elevated and MDSC compartment is altered in CCN4 KO compared to WT Py8119 tumors. In a size-matched experimental design, the tumor-infiltrating lymphocytes were quantified by flow cytometry using antibody panels focused on the T cell (A, B, and C) and the myeloid (D, E, and F) compartments. Mice received either WT Py8119 cells (black circles) or one of two independently generated CCN4 KO variants (red + and blue x, n = 5 for all groups). Results highlight the CD45+ fraction among total Live events measured using the T cell (A) and myeloid (D) panels. The proportion of infiltrating CD4+ T cells within the live CD45+ compartment (B) was not different but the CD25+ fraction of live CD4+ T cells (C) was increased in CCN4 KO tumors. The MDSC compartment was shifted from PMN-MDSC (E, Live CD45+ CD11b+ Ly6G+ Ly6C^*int*^ events) to Mo-MDSC (F, Live CD45+ CD11b+ Ly6G-Ly6C+ events) upon CCN4 KO. p-values calculated between WT and pooled CCN4 KOs using two-sided Student’s t test.

Similar to the Py230 experiments, the myeloid compartment also underwent a compositional shift: PMN-MDSCs were significantly reduced (p-value *<* 0.01, Figure 4E) while Mo-MDSCs increased (p-value *<* 0.02, Figure 4F) in CCN4 KO tumors. This shift is functionally significant, as PMN-MDSCs are generally considered more potently immunosuppressive than Mo-MDSCs and are associated with poor prognosis in multiple cancer types. Moreover, we observed a similar shift in the myeloid compartment from PMN-MDSCs to Mo-MDSCs upon CCN4 KO using the YUMM1.7 model, a transplantable mouse model of melanoma, and that this phenotype can be rescued by inducing CCN4 expression in KO cells (6). Given that our primary endpoint - tumor growth in immunocompetent mice - showed significant CCN4-dependent differences, and given that CD8+ T cells were clearly elevated in CCN4 KO tumors, we interpret the overall immune phenotype upon CCN4 KO as favorable for anti-tumor immunity.

### CCN4 regulates a conserved immunosuppressive secretome

To identify potential mechanisms by which CCN4 promotes immunosuppression, we profiled the secretomes of WT and CCN4 KO cell lines using a multiplexed antibody array that detects 111 cytokines, chemokines, and growth factors. CCN4 knockout substantially altered the secretory profile in both Py230 and Py8119 cells (Figure 5A, B). Several secreted factors with known immunosuppressive or protumorigenic functions were downregulated upon CCN4 loss, including Serpin E1 (also known as PAI-1), which promotes MDSC accumulation (12), and CCL20, which recruits regulatory T cells (13), in the Py230 model and CXCL1, which attracts immunosuppressive neutrophils (14), in the Py8119 model. Conversely, some factors were upregulated upon CCN4 loss, including CXCL10, a T cell chemoattractant associated with improved anti-tumor immunity (15), in the Py8119 model.

**Fig. 5.**
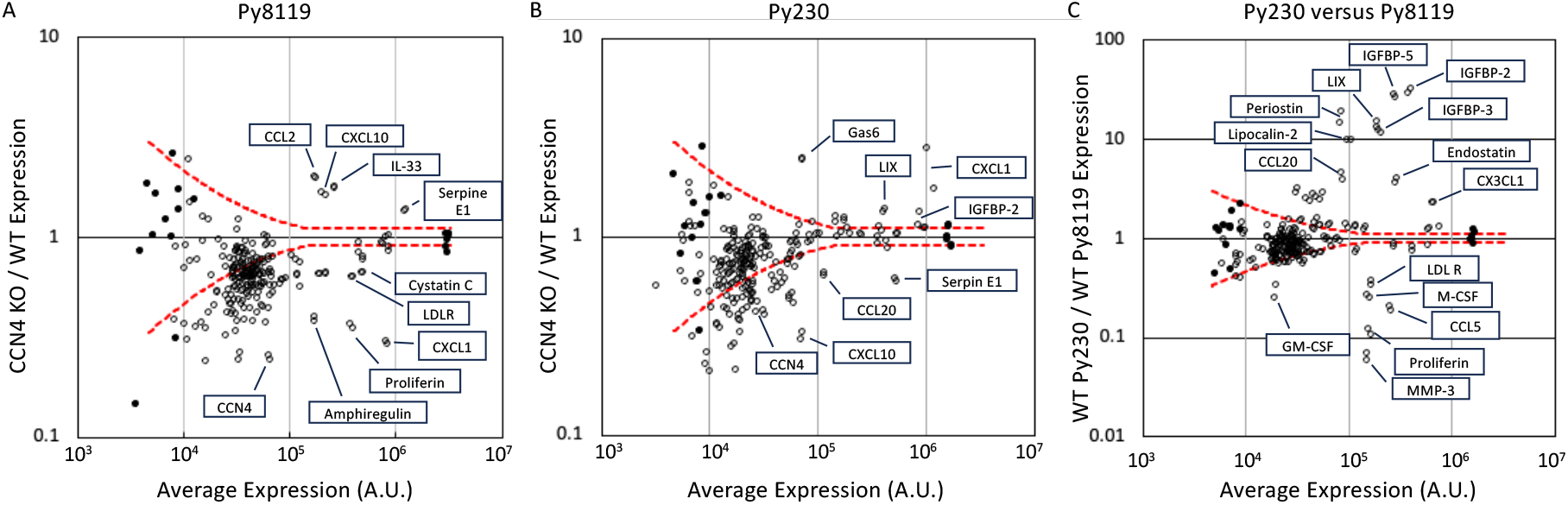
The profile of cytokines, chemokines, and growth factors secreted into media conditioned by Py230 and Py8119 cells are changed upon CCN4 KO. Results from R&D Systems’ Mouse XL Cytokine Array Kit of cytokine, chemokine, and growth factor expression by WT versus CCN4 KO cells derived from Py8119 (A) and Py230 (B) lines *in vitro*. (C) Compares the WT profiles between the Py8119 and Py230 cell lines. In presenting the combined results for assaying each cell line’s secretome using a single biological replicate, open circles represent results for specific cytokine probes, which are spotted in duplicate on the array, and filled circles represent positive and negative controls. Dotted red lines indicate Z-scores of 3 and −3. Particular secreted factors that are differentially expressed are annotated with their respective names.

As the Py230 and Py8119 cell lines represent epithelial-like and mesenchymal-like adenocarcinoma, it is not surprising that the baseline secretomes of Py230 and Py8119 cells differed substantially from one another (Figure 5C) and reflect their distinct transcriptional programs and differentiation states. It was interesting, though, that the differential changes induced by CCN4 knockout were remarkably consistent. Moreover, principal component (PC) analysis across four different cancer cell lines − the two breast cancer models (Py230 and Py8119) and two melanoma models (B16F0 and YUMM1.7) − revealed that CCN4 knockout induced similar secretome shifts across all four lines (Figure 6). PC1, which explained 99% of the variance, separated WT from CCN4 KO variants regardless of cell line origin. As the majority of proteins had normally distributed loading scores centered at −0.091, the protein names with loading coefficients for PC1 below −0.0925 identify key proteins that are downregulated upon CCN4 KO across different cancer types (Supplemental Table S1). As there are no common proteins that are upregulated upon CCN4 KO, specific observations depend on the particular cell model and that CCN4 generally enables rather than represses the expression of proteins.

**Fig. 6.**
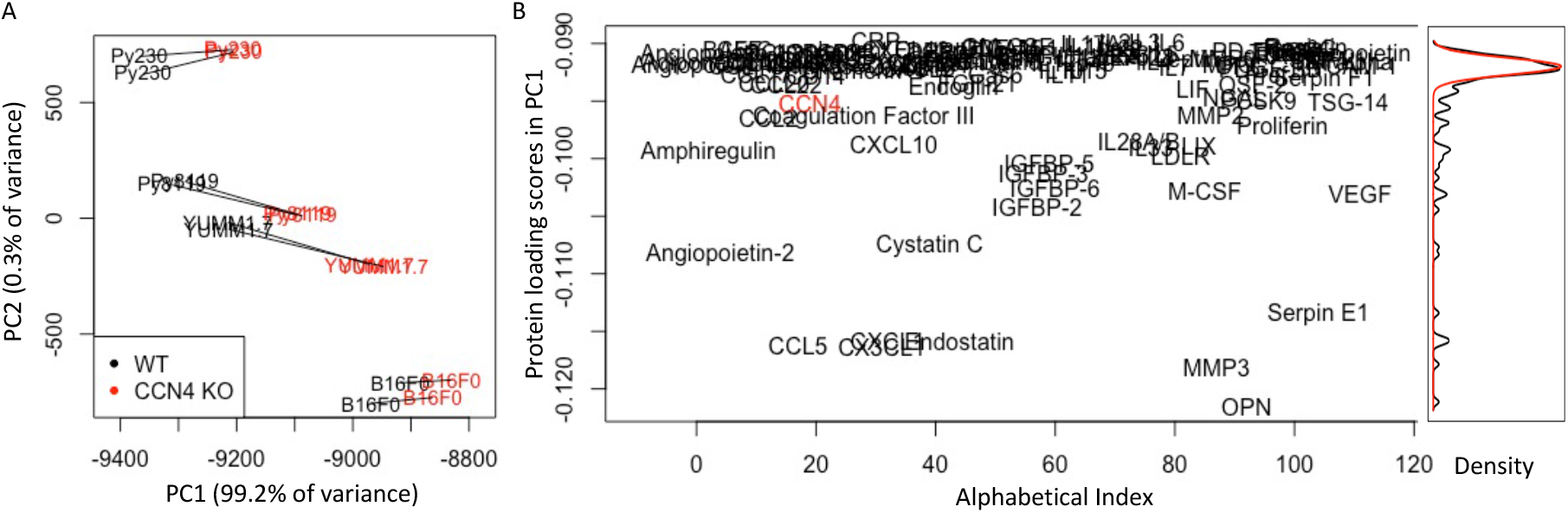
While the profile of cytokines, chemokines, and growth factors secreted into media conditioned by WT Py230, Py8119, B16F0 and YUMM1.7 cells are different, the changes in profiles associated with CCN4 KO are similar. (A) Projections of WT versus CCN4 KO variants along PC1 and PC2 axes for B16F0, YUMM1.7, Py230, and Py8119 cell lines. The location of duplicates of each cell line label correspond to technical replicates on the array. The percentage of variance explained by each principal component is indicated in the axis label. (B) Projection of the measured proteins based on the loading scores associated with PC1 (y-axis) listed from left to right in alphabetical order. The curves on the right represent the density distribution of the PC1 loading scores (black) and a normal distribution centered at −0.091 with a standard deviation of 0.00074 (red).

### Genetic alterations promote CCN4 expression through beta-catenin pathway activation

While we had previously reported the near-universal upregulation of *CCN4* gene expression in primary tumor samples obtained from patients diagnosed with invasive breast cancer and that this increase in *CCN4* gene expression correlated with a decrease in type 1 cell-mediated cytotoxic immunity (3), genetic drivers underlying this upregulation remained unclear. In connecting elevated CCN4 expression with disruption of adherens junctions that occurs during invasion, we identified an interlocked positive and negative feedback network motif ((16) and summarized in Figure 7A, B) where genetic alterations lead to increased CCN4 expression via constitutive activation of the Wnt pathway.

**Fig. 7.**
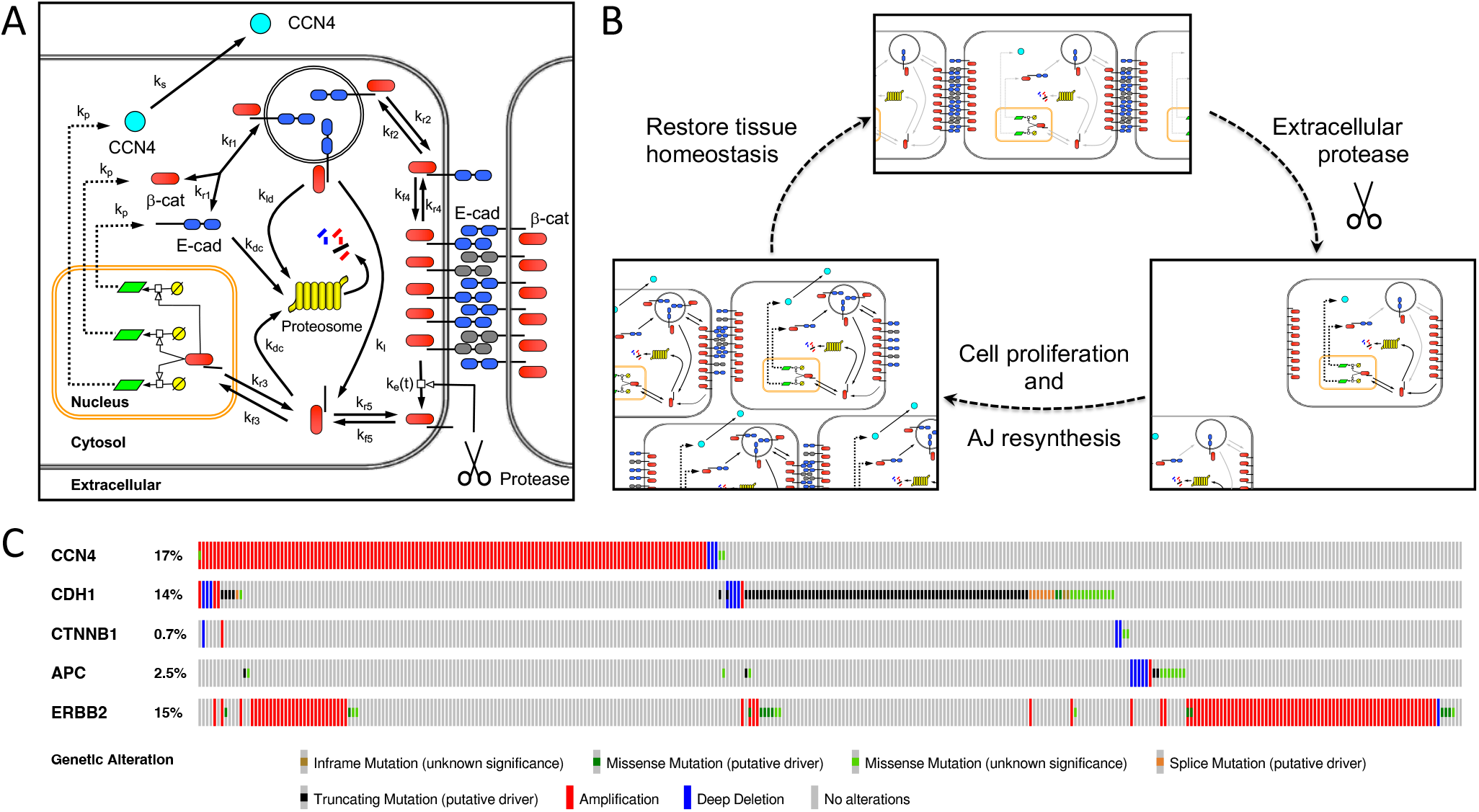
Schematic diagram of the signaling network associated with non-canonical activation of beta-catenin upon disrupting adherens junctions and associated mutational landscape observed in invasive breast cancer. (A) Schematic diagram of the network topology associated with the adherens pathway activity of beta-catenin. Adherens junctions are formed by homotypic interactions between the extracellular cadherin domains of a single pass transmembrane protein E-cadherin (E-cad: blue double oval). The cytoplasmic tail of E-cadherin provides a scaffold for a multi-protein complex that includes beta-catenin (*β*-cat: red oval). (B) Upon the proteolytic cleavage of adherens junctions, a cytoplasmic fragment of E-cadherin and associated catenins (tBE) are transported to the cytoplasm. In the cytoplasm, the cytoplasmic fragment of E-cadherin and associated catenins can either enter the nucleus or undergo proteasomal degradation. In the nucleus, this multi-protein complex promotes the transcription and translation of CCN4, beta-catenin, and E-cadherin, among other factors. mRNA is represented by light green parallelogram. Once synthesized, CCN4 (light blue circle) is secreted. Newly synthesized beta-catenin and E-cadherin reform the multi-protein complex and are transported to the cell membrane to re-establish adherens junctions. If no E-cadherin binding sites are present, the multi-protein complex is internalized and degraded. (C) Graphical summary of mutations that include both changes in single nucleotides and in copy numbers for breast invasive carcinoma (TCGA, Cell 2015) samples. Of the 816 samples contained in the dataset obtained through cBioPortal (retrieved 11/10/2025), the queried genes are altered in 336 (41%) of the samples.

To determine whether genetic alterations supporting this mechanism are enriched in breast cancer, we analyzed The Cancer Genome Atlas (TCGA) breast invasive carcinoma dataset (Figure 7C). Interestingly, genetic alterations that activate this pathway were enriched in this invasive breast cancer cohort. For instance, copy number amplifications of CCN4, which would increase the amount of mRNA produced in response to a transcriptional cue, is enriched in 17% of samples. As a point of reference, the most prevalent genetic alteration involving *ERBB2* is copy number amplifications at a frequency of 15%. Mutations in *CDH1* gene are present in 14% of samples, where the majority of missense or truncating mutations in *CDH1* occur in the extracellular cadherin domains. These mutations in the extracellular domains have the potential to disrupt homotypic binding and promote constitutive pathway activity (17). In this invasive breast cancer dataset, mutations in *CTNNB1* are not prevalent as they occur with a frequency of less than 1%. In other cancers, Exon 3, which is in the N-terminus and helps target the protein to the proteosome for degradation, preferentially harbors mutations that increase nuclear accumulation of beta-catenin (18). At a prevalence of 3%, mutations in *APC* that lead to impaired function were also enriched. As APC negatively regulates beta-catenin activity, these mutations would also lead to enhanced pathway activity (19). Given that mutations in these four genes mentioned all have the same functional response, it is interesting to note that these mutations are largely non-overlapping, suggesting that tumors are under selective pressure to activate this pathway but that multiple genetic routes can achieve this outcome.

The observation that CCN4 genomic amplification occurs at a frequency comparable to ERBB2 amplification, combined with the functional non-overlap of mutations affecting the beta-catenin pathway, suggests that CCN4-mediated immunosuppression may be under strong selective pressure during tumor evolution. This genomic analysis provides a mechanistic link between adherens junction disruption during invasion, constitutive beta-catenin signaling, CCN4 upregulation, and the immunosuppressive tumor microenvironment observed in our experimental models.

### CCN4 as a therapeutic target in breast cancer

Our findings establish CCN4 as a driver of tumor-induced immunosuppression in breast cancer, extending previous observations in melanoma to a second major cancer type. The immune-dependent growth phenotype, reduced splenomegaly, enhanced T cell infiltration, favorable remodeling of the MDSC compartment, and conserved secretome changes collectively demonstrate that CCN4 actively shapes the tumor immune microenvironment to suppress anti-tumor immunity.

These intriguing results motivate several mechanistic follow-on questions. For instance, the precise cellular targets of CCN4 within the tumor microenvironment remain to be definitively established. CCN4 is a secreted matricellular protein that can interact with multiple cell surface receptors, including integrins and receptor tyrosine kinases. While identifying a defined cell surface receptor might be a first-pass thought, the unsuccessful focus of the scientific community on identifying the “CCN4 cell surface receptor” was a barrier for progress within the field (20). Instead, CCN proteins should be thought of as organizers of a bidirectional interaction network between a cell and the extracellular matrix by docking extracellular proteins with cell surface proteins like integrins and can exhibit both autocrine and paracrine action. Previously we have shown that CCN4 can act in a paracrine fashion by directly inhibiting T cell response to IL-12 (21) and CD8+ T cell function (6). IL-12 is an important secreted signal that connects the myeloid compartment, like dendritic cells, with engaging the cytotoxic effector function, like CD8+ T and NK cells (22, 23). Tonic IL-12 signaling within the tumor microenvironment underpins the enhanced anti-tumor immune response engaged by immune checkpoint blockade (24). The global changes in secretome profiles exhibited upon CCN4 KO suggest that CCN4 acts in an autocrine fashion on these cancer cells. This autocrine action complicates interpreting how CCN4 KO impacts the myeloid compartment, as changes in MDSC-attracting chemokines, like CXCL1 and CCL20, provide an indirect mechanism for the observed phenotype. Characterizing CCN4 action in a well-defined cellular context may help clarify how CCN4 orchestrates a cellular response.

These results have important therapeutic implications. The upregulation of CCN4 in essentially all invasive breast cancers, combined with its correlation with poor outcomes and dampened anti-tumor immunity in clinical datasets, positions CCN4 as an attractive therapeutic target. Importantly, because CCN4 functions as a secreted matricellular protein rather than a cell-intrinsic oncogene, it may be more readily targeted with antibody-based therapeutics or small molecule inhibitors compared to intracellular targets. Given the autocrine and paracrine nature of CCN4 action, understanding how a particular therapeutic strategy parses autocrine versus paracrine action may have a significant influence on therapeutic efficacy. The observation that CCN4 knockout shifts the tumor immune microenvironment toward a more immunologically “hot” phenotype − with increased T cell infiltration and reduced immunosuppressive myeloid cells − suggests that CCN4 blockade could synergize with checkpoint immunotherapies. Patients whose tumors fail to respond to anti-PD-1 or anti-CTLA-4 antibodies often exhibit poor T cell infiltration and accumulation of immunosuppressive myeloid cells, precisely the features that CCN4 promotes. Thus, combining CCN4 inhibition with checkpoint blockade represents a rational strategy to overcome resistance and expand the population of breast cancer patients who benefit from immunotherapy.

The conservation of CCN4’s immunosuppressive program across breast cancer and melanoma further supports its potential as a broadly applicable therapeutic target. The similar secretome changes observed across four independent cell lines from two distinct cancer types suggest that CCN4 regulates fundamental aspects of tumor-immune crosstalk that transcend tissue-specific contexts. This conservation increases the likelihood that therapeutic strategies targeting CCN4 could be effective across multiple cancer indications.

Future studies should focus on several key areas to advance CCN4 as a therapeutic target. First, the specific immune cell populations that directly respond to CCN4 signaling need to be identified through receptor expression profiling and functional studies. Second, the intracellular signaling pathways downstream of CCN4 that drive immunosuppressive programs should be elucidated to identify additional druggable nodes. Third, the development and preclinical evaluation of CCN4-blocking antibodies or small molecule inhibitors in combination with standard immunotherapies will be essential. A related point is that an acute tumor challenge using a KO line, like this study, does not address whether CCN4 inhibition can reverse established immunosuppression, which is a critical translational question. Finally, clinical validation studies examining CCN4 expression, copy number status, and immune infiltration patterns in patient cohorts treated with immunotherapy will help refine patient selection strategies and establish CCN4 as a predictive biomarker for therapeutic response. Patients whose tumors harbor CCN4 amplifications may represent a particularly relevant population for CCN4-targeted therapies, as these tumors may be especially dependent on CCN4-mediated immune evasion.

## Methods

### Reagents and Cell culture

All biochemical reagents were obtained from commercial sources and used according to the suppliers’ recommendations unless otherwise indicated. Mouse breast cancer Py230 (Purchased June 2017, CRL-3279) and Py8119 (Purchased October 2018, CRL-3278) cell lines were purchased from American Type Culture Collection (ATCC) and grown as recommended from ATCC. The cells were maintained in F-12K Medium (Cellgro/Corning, NY) supplemented with 5% heat-inactivated FBS (FBS, Hyclone, UT), 0.1% MITO+ Serum Extender (Corning, NY), L-glutamine (Lonza, NJ) and 1% penicillin-streptomycin (Gibco, ThermoFisher Scientific, MA) at 37^*o*^C in 5% CO2. All cells lines were revived from frozen stock, used within 10-15 passages that did not exceed a period of 6 months, and routinely tested for mycoplasma contamination by PCR.

### Creation of CCN4-Knockout Cells Using CRISPR/Cas9 System

To achieve high specificity and reduce variability in genetic backgrounds, CRISPR/Double Nickase systems were selected to knock out the Ccn4 gene. Two pairs of mouse Ccn4 Double Nickase Plasmids (sc-423705-NIC and sc- 423705-NIC-2), targeting the mouse Ccn4 gene at different locations, were purchased from Santa Cruz Biotechnology (Dallas, Texas) and used in Py230 and Py8119 cells. Following manufacturer’s instructions, cells were transfected with individual set of plasmids, which also express a puromycin-resistance gene. Cells were selected by puromycin for five days to achieve 100% transfection efficiency. Surviving cells were counted and plated into 96-well plate with a density of cell/well. After one week, single clones were isolated and expanded on 6-well plates. The cell culture media from those wells were used for CCN4 ELISA to characterize knockout clones. The identified CCN4-knockout cells were further expanded and used for subsequent experiments.

### In vitro studies

To measure CCN4 secretion from each WT cell line and KO variant, cells were grown for 48 hours to reach about 80% confluence and the media was filtered for ELISA (enzyme-linked immunosorbent assay) analysis using a Mouse CCN4 Quantikine ELISA Development Kit (R&D Systems, Minneapolis, MN). To collect tumor-conditioned media (TCM), wild-type and CCN4 KO variants of Py230 and Py8119 cells were grown in complete F-12K medium until 80% confluency, washed with PBS (Cellgro/Corning, NY) and incubated for 48 h in FBS-free F-12K. TCM were then centrifuged at 3,000 g and 4^*o*^C for 15 min, and the supernatant collected and filtered. Cytokines, chemokines, and growth factors in TCM were detected with the Proteome Profiler Mouse XL Cytokine Array (R&D Systems, MN), following the manufacturer’s instructions. As each antibody was spotted in duplicate on the array, intensity values for each spot were considered a technical replicate and used in subsequent analysis. Raw intensity results from the scanned arrays were normalized to give similar dynamic ranges for the positive and negative controls. Normalized intensity values are provided in Supplemental File 1: Antibody Array Summary.

### Mice and in vivo tumor studies

For assaying in vivo tumor growth, C57BL/6Ncrl and NOD.Cg-Prkdc^*scid*^ Il2rg^*tm*1*Wjl*^/SzJ (NSG) mice (6-8 weeks of age, female) were purchased from Charles River Laboratories and The Jackson Laboratory, respectively. Mice were housed in sterilized microisolator cages in the university vivarium, and facility sentinel animals were regularly screened for specific pathogen agents. Mice were randomly assigned to treatment groups and co-housed following tumor initiation. Animals were housed with a 12-hour light/dark cycle (light 6 am to 6pm), temperature nominally 74 degrees F, and humidity 50%. All animal experiments were approved by the West Virginia University (WVU) Institutional Animal Care and Use Committee and performed on-site (IACUC Protocol #1604002138). No animals were excluded from reporting of the study with n ≥5 in each experimental group. Results representative of one of two independent experiments. As is custom in the field, investigators were not blinded to cohort membership when measuring tumor size. For tumor challenge, mice were implanted subcutaneously (s.c) on the right flank or orthotopically injected into the #4 mammary fat pad with 2.5 *×* 106 Py230 (WT and KO variants) or 5 *×* 105 Py8119 (WT and KO variants) cells in PBS. Once palpable, tumor growth was monitored every two days by measuring two perpendicular diameters of the tumor using calipers. The tumor volume was calculated according to the formulation (*π*/6 *×* length *×* width^2^), where the width is the smaller dimension of the tumor. For ethical considerations, mice were sacrificed when tumor burden was significant enough to effect daily life (difficulty moving, weight loss, inability to feed or gain access to water, etc). The cohort size was not prespecified as this was an exploratory study and not designed to test for a pre-specified effect.

### Flow cytometry

To assay tumor infiltrating lymphocytes once tumors from both WT and CCN4 KO cohorts reached a similar size, the tumors were surgically removed from mice in both arms of the study (WT and CCN4 KO) after euthanasia and processed into single cell suspensions. This nominally occurred at Day 30 with the WT Py230 cells and at Day 57 with the CCN4 KO1 variant reaching 300 mm^3^ in size. For mice receiving Py8119 variants, the experiment was stopped when tumors reached 1000 mm^3^ in size, which nominally occurred at Day 30 with WT cells, at Day 36 with CCN4 KO1 variant, and Day 44 with CCN4 KO2 variant. Five tumors were processed separately for each WT and CCN4 KO variants. Single-cell suspensions were obtained by enzymatically digesting the excised tumors using the Tumor Dissociation Kit and gentleMACS C system (Miltenyi Biotec, Auburn, CA). In addition to following the manufacturer’s instructions, the gentleMACS program 37C_m_TDK_2 was used for Py230 tumors. Following lysing of the red blood cells, the remaining singlecell suspensions were washed and stained with Live/Dead Fixable Red Dead Cell Stain Kit (ThermoFisher). Following blocking with Mouse BD Fc Block (purified rat antimouse CD16/CD32 antibodies, BD Biosciences), the surface of the cells obtained from the Py230 cohorts were stained with a single spectral antibody mix that focused on T cells (CD45, CD3, CD4, CD8a, CD25, and FoxP3), NK and B cells (CD45, CD3, B220, and NK11), and myeloid cells (CD45, CD11b, CD11c, Ly-6C, Ly-6G, F4/80, and MHCII) and quantified by flow cytometry. The specific antibodies and dilutions used are listed in Supplementary Table S1. Events were acquired using a Cytek Aurora (Cytek) spectral flow cytometer with SpectroFlo software, where the fluorescence intensity for 38 channels distributed across three laser excitation beams plus forward and side scatter intensities were reported as a pulse area with 18-bit resolution. Flow cytometric data were exported as FCS3.0 files and analyzed with a custom R script. Details of the processing steps are described in Supplemental methods.

Following blocking with Mouse BD Fc Block (purified rat anti-mouse CD16/CD32 antibodies, BD Biosciences), the surface of the cells obtained from the Py8119 cohorts were stained with one of two different antibody mixes that focused on T, NK, and B cells (CD45, CD3, CD4, CD8, B220, NK11, and CD25), and myeloid cells (CD45, CD11b, CD11c, Ly-6C, Ly-6G, F4/80, and MHCII) and quantified by flow cytometry using a BD LSRFortessa and FACSDiva software (V8.0 BD Biosciences). Unstained samples were used as negative flow cytometry controls. Single-stain controls were used to establish fluorescence compensation parameters. For TIL analysis, greater than 2 *×* 105 events were acquired in each antibody panel in each biological replicate. Flow cytometric data, where the fluorescence intensity for each parameter was reported as a pulse area with 18-bit resolution, were exported as FCS3.0 files and analyzed using R/Bioconductor (V3.6.1), as described previously (25). The typical gating strategies for T cells, NK and B cells, and myeloid cells are shown in Supplementary Figures S2 and S3, respectively. The statistical difference in tumor infiltrating lymphocytes between WT and CCN4 KO variants was assessed using log-transformed values and a two-tailed homoscedastic Student’s t test. A p-value of less than 0.05 was considered statistically significant.

### Statistical analysis

Results from all univariate experiments were analyzed by a two-tailed, unpaired Student’s t-test. To estimate cumulative survival probability, KaplanMeier survival curves were estimated from the cohort time-to-event data, where a presence of a tumor greater than 100 mm^3^ was considered the endpoint event. Statistical significance associated with a difference in time-to-event between groups was estimated using the Peto & Peto modification of the Gehan-Wilcoxon test, as implemented in the “survival” (V3.6-4), “survminer” (V0.5.0), and “stats” (V4.1.1) packages in R. A p-value of *<*0.05 was considered statistically significant. Unless otherwise specified, quantitative results were summarized as mean *±* standard deviation (s.d.) and overlaid on individual results. In analyzing the Mouse XL Cytokine Array, positive and negative controls were used to to establish a null distribution, that is that an observed difference in abundance is explained by random chance. The curves enclosing 68% of the null distribution are calculated by regressing a curve to the standard deviation of the observed replicates of an antibody probe on an array among a subset of samples of a similar log abundance versus the average of the sample subset. Statistical significance associated with differential expression can be estimated by comparing the vertical distance of a measured secreted factor from the x-axis relative to the vertical distance between the 68th percentile null distribution curve and the x-axis. The comparison of these two vertical distances is proportional to a z-score. Observations that lie outside of the 99.7% null distribution curves, that is Z-scores greater than 3 and less than −3, are considered significant and merit further consideration. By regressing an additive-multiplicative error model to the probe intensities on the array, the raw intensity values were transformed using a variance stabilizing model as described in (26). These variance stabilized intensity values were used for principal component analysis as implemented in the “stats” (V4.1.1) package in R.

## ACKNOWLEDGEMENTS

This work was supported by National Science Foundation (NSF CBET-1644932 to DJK) and National Cancer Institute (NCI 1R01CA193473 to DJK). The content is solely the responsibility of the authors and does not necessarily represent the official views of the NSF or NCI. The authors would like to thank Audry Fernandez and the WVU Flow Cytometry and Single Cell Analysis Core for their assistance in conducting this study.

## AUTHOR CONTRIBUTIONS

Conceptualization: DJK; Study Design: DJK; Data Acquisition: DJK, WD, AR, AP; Data Analysis: DJK, AP; Data Interpretation: DJK; Funding acquisition: DJK; Methodology: DJK; Project administration: DJK; Software: DJK; Supervision: DJK; Writing − original draft: DJK, WD, AR, AP; Writing − review & editing: DJK.

## Supplementary Information

This supplemental file contains:

**Fig. S1.**
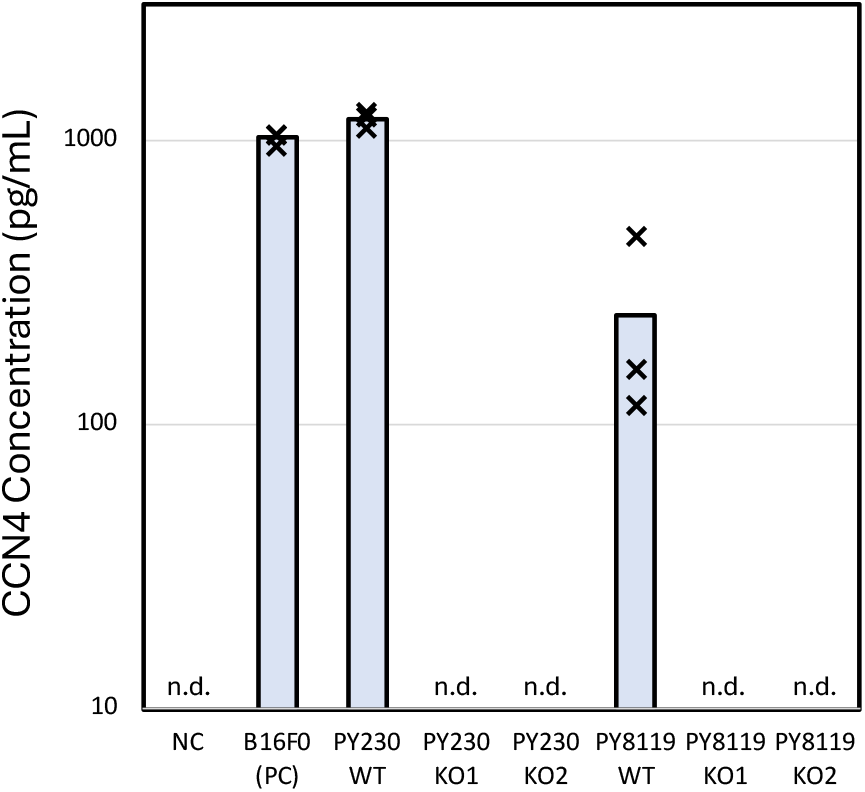
Validation of CCN4 KO in Py230 and Py8119 cells. CCN4 concentration as quantified by ELISA in media conditioned for 48 hours by WT Py230 and Py8119 cells and two independent CCN4 KO variants for each parental cell line. Cell-free media was used as a negative control and media conditioned by B16F0 cells was used as a positive control. Individual data points represent technical replicates and values reported below quantification threshold are indicated as n.d.

**Fig. S2.**
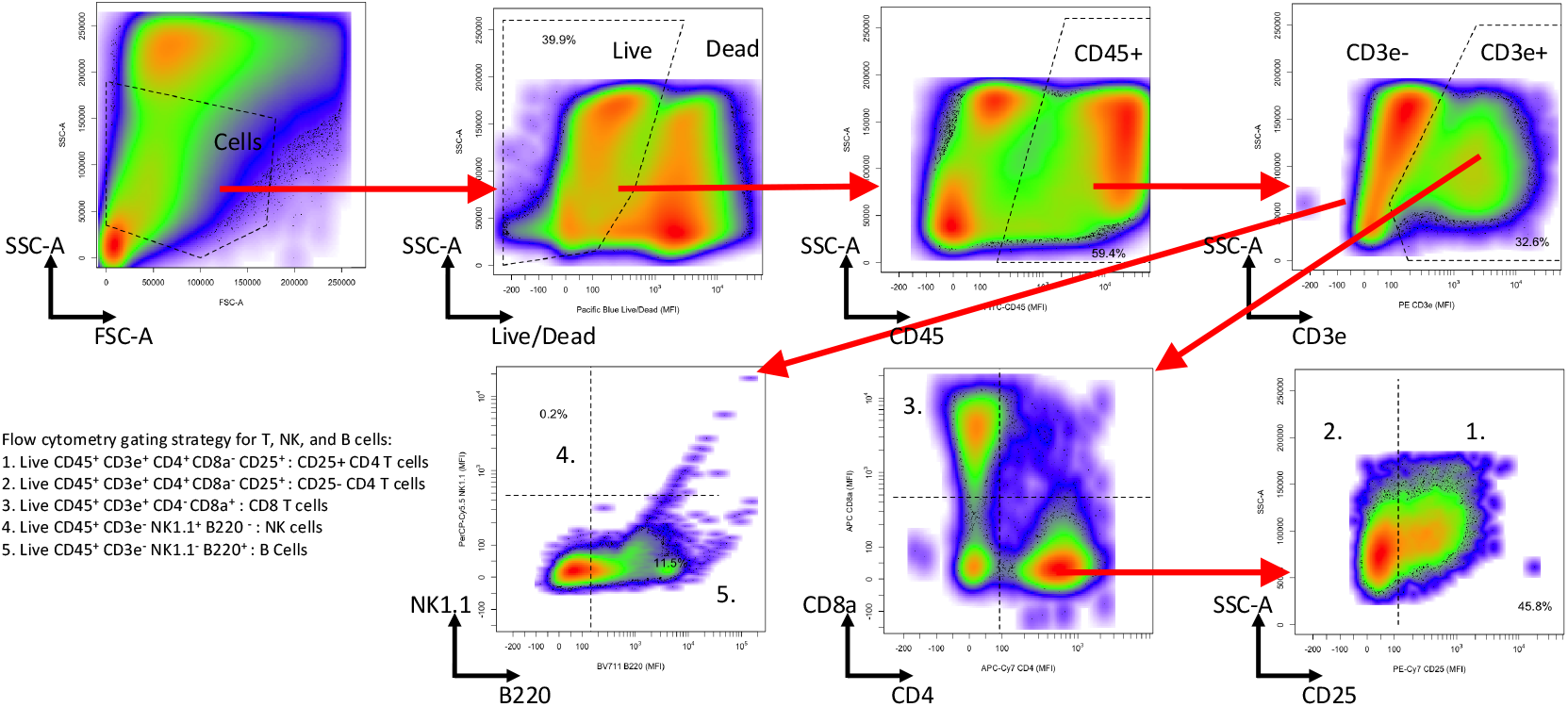
Flow cytometry gating strategy for T, NK, and B cells. After gating on cells, Live Dead Pacific Blue staining versus side scatter area was used to then gate for Live cells. Then CD45 staining versus side scatter area was used to gate for Live CD45^+^ cells. Live CD45^+^ events were then gated based on CD3e^+^ expression. Live CD45^+^ CD3e^+^ cells were further subdivided into CD8^+^ T cells (live CD8a^+^ CD4^−^ CD3e^+^ CD45^+^ cells) and CD4 T cells (live CD4^+^ CD8a^−^ CD3e^+^ CD45^+^ cells). CD4 T cells were further gated based on CD25 expression into high and low expression subsets. Live CD45^+^ CD3e^−^ events were subdivided into B cells (live NK1.1^−^ B220^+^ CD3e^−^ CD45^+^ cells) and NK cells (live NK1.1^+^ CD3e^−^ CD45^+^ cells).

**Fig. S3.**
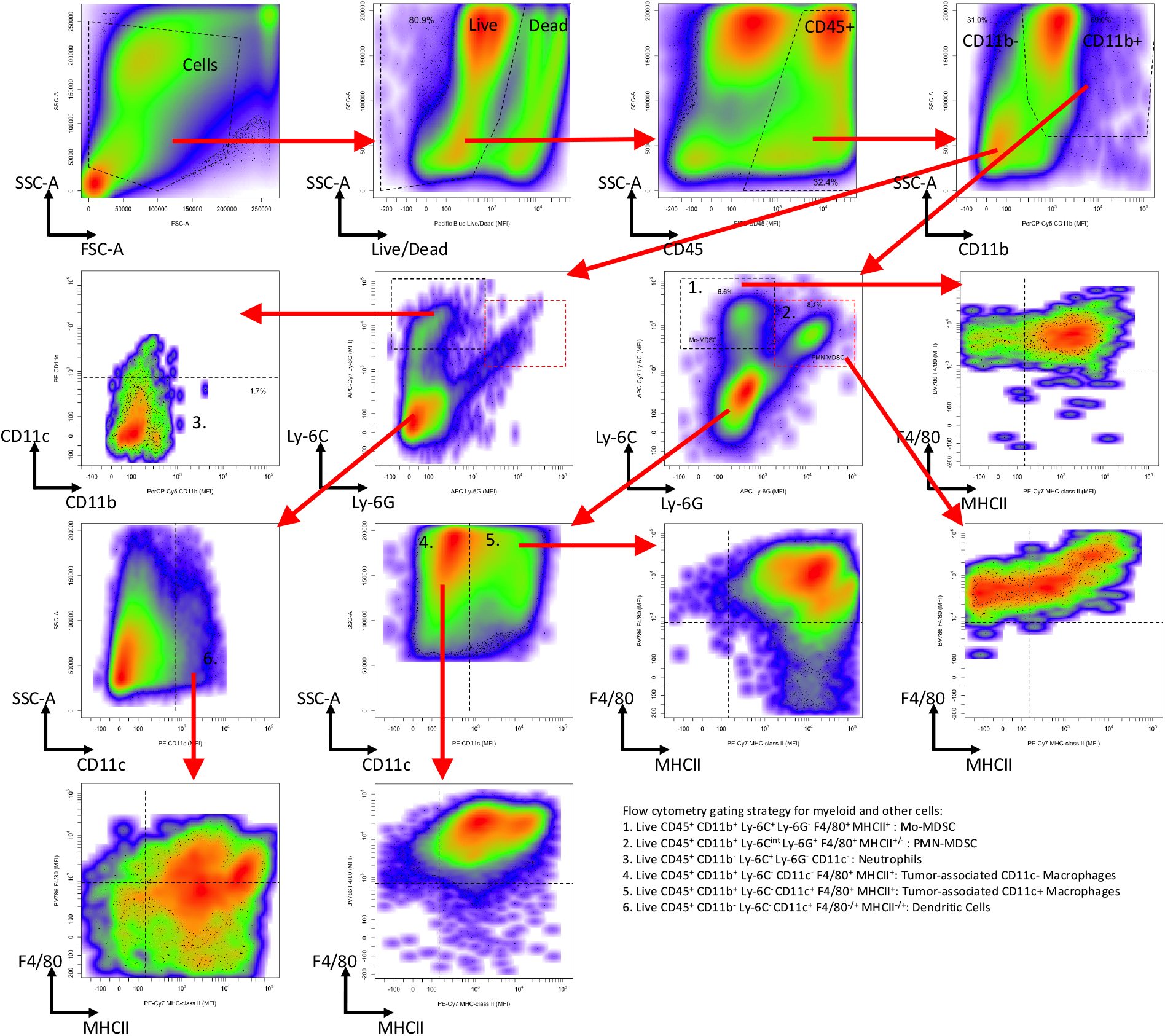
Flow cytometry gating strategy for Tumor associated neutrophils (TAN) and myeloid cell subsets. After gating on cells, Live Dead Pacific Blue staining versus side scatter area was used to then gate for Live cells. Then CD45 staining versus side scatter area was used to gate for Live CD45^+^ cells. Live CD45^+^ events were then subdivided into subsets based on CD11b staining followed by Ly-6C versus Ly-6G staining. From the CD11b^+^ gate, myeloid-derived suppressor cells (MDSC) were subdivided into Monocytic-MDSC (Mo-MDSC: live CD45^+^ CD11b^+^ Ly-6C^+^ Ly-6G^−^ cells) and Polymorphonuclear-MDSC (PMN-MDSC: live CD45^+^ CD11b^+^ Ly-6C^+^ Ly-6G^−^). In addition, cells from the CD11b^+^ gate that were not Mo-MDSC or PMN-MDSC were gated based on CD11c expression into tumor-associated CD11c^−^ and CD11c^+^ macrophages. The CD11b^−^ subset included tumor-associated neutrophils (TAN) (live CD45^+^ CD11b^−^ Ly-6C^+^ Ly-6G^−^ CD11c^−^) and dendritic cells (live CD45^+^ CD11b^−^ Ly-6C^−^ CD11c^+^).

**Table S1.** Top 40 proteins based on PC1 loading scores.

| Protein | Loading Score | Protein | Loading Score |
| --- | --- | --- | --- |
| OPN | -0.12156 | IL28A/B | -0.09849 |
| MMP3 | -0.11817 | Proliferin | -0.09709 |
| CX3CL1 | -0.11631 | CCL2 | -0.09642 |
| CCL5 | -0.11616 | Coagulation Factor III | -0.09642 |
| Endostatin | -0.11590 | MMP2 | -0.09625 |
| CXCL1 | -0.11578 | CCN4 | -0.09517 |
| Serpin E1 | -0.11353 | TSG-14 | -0.09506 |
| Angiopoietin-2 | -0.10828 | PCSK9 | -0.09498 |
| Cystatin C | -0.10756 | NGAL | -0.09466 |
| IGFBP-2 | -0.10422 | LIF | -0.09396 |
| VEGF | -0.10301 | Endoglin | -0.09385 |
| M-CSF | -0.10284 | OSF-2 | -0.09358 |
| IGFBP-6 | -0.10258 | CCL22 | -0.09350 |
| IGFBP-3 | -0.10133 | CCL20 | -0.09348 |
| IGFBP-5 | -0.10036 | FGF-21 | -0.09335 |
| LDLR | -0.09975 | Serpin F1 | -0.09310 |
| Amphiregulin | -0.09944 | CD14 | -0.09273 |
| IL33 | -0.09899 | CCL17 | -0.09271 |
| LIX | -0.09885 | IL11 | -0.09269 |
| CXCL10 | -0.09870 | Gas6 | -0.09263 |

**Table S2.**
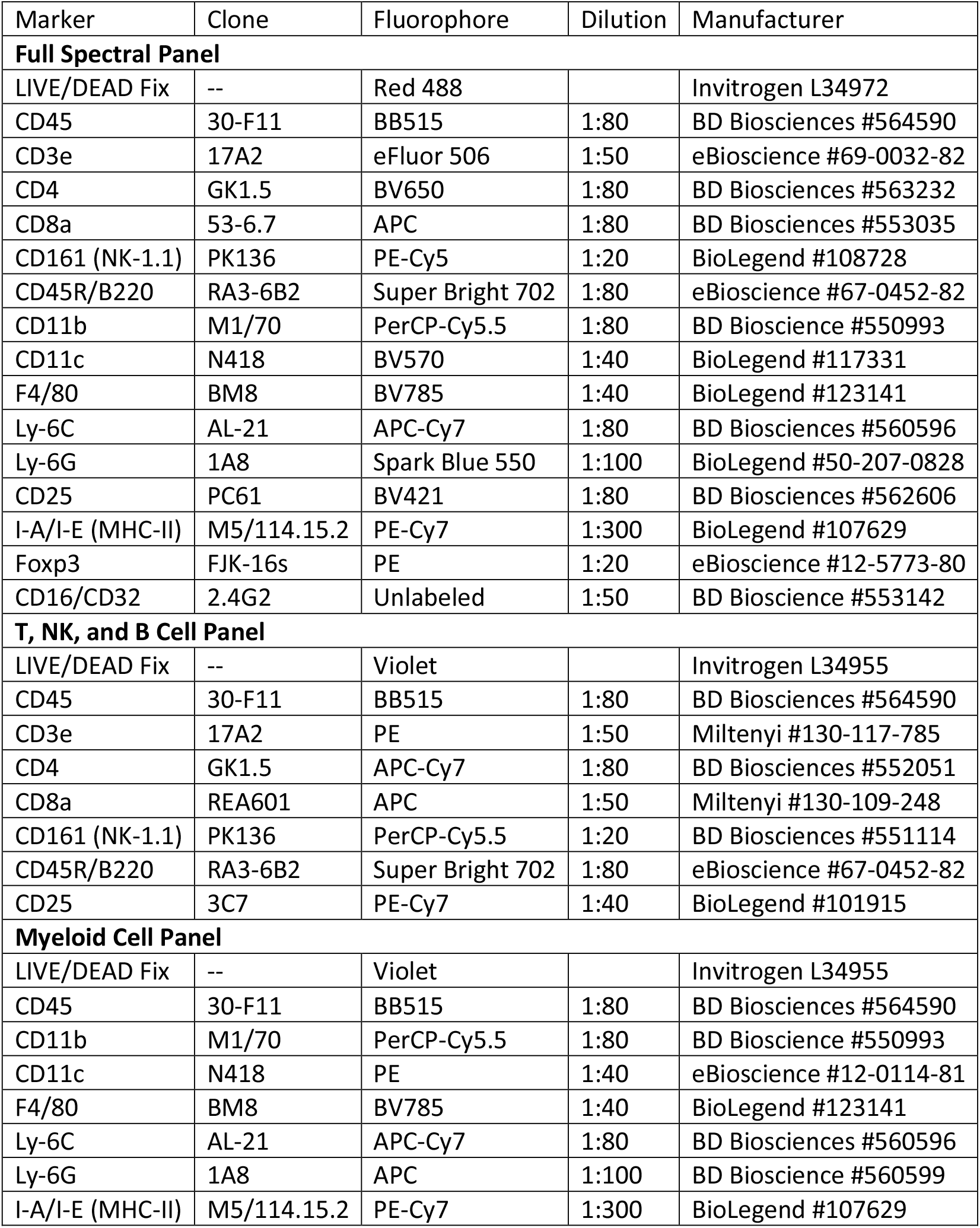
List of fluorophore-conjugated anti-mouse antibodies using to quantify cell subsets by flow cytometry.

## Bibliography

1. D. J. Klinke. Is immune checkpoint modulation a potential therapeutic option in triple negative breast cancer? Breast Cancer Res, 16(1):457, Nov 2014.

2. J. Cortes, D.W. Cescon, H.S. Rugo, Z. Nowecki, S. Im, M.M. Yusof, C. Gallardo, O. Lipatov, and C.H. et al. Barrios. Pembrolizumab plus chemotherapy versus placebo plus chemotherapy for previously untreated locally recurrent inoperable or metastatic triple-negative breast cancer (KEYNOTE-355): a randomised, placebo-controlled, double-blind, phase 3 clinical trial. Lancet, 396:1817–1828, 2020.

3. D. J. Klinke. Induction of Wnt-inducible signaling protein-1 correlates with invasive breast cancer oncogenesis and reduced type 1 cell-mediated cytotoxic immunity: a retrospective study. PLoS Comput. Biol., 10(1):e1003409, Jan 2014.

4. Wentao Deng, Audry Fernandez, Sarah L McLaughlin, and David J Klinke. WNT1-inducible signaling pathway protein 1 (WISP1/CCN4) stimulates melanoma invasion and metastasis by promoting the epithelial-mesenchymal transition. J Biol Chem, 294(14):5261–5280, 2019.

5. W. Deng, A. Fernandez, S. L. McLaughlin, and D. J. Klinke. Cell Communication Network Factor 4 (CCN4/WISP1) Shifts Melanoma Cells from a Fragile Proliferative State to a Resilient Metastatic State. Cell Mol Bioeng, 13(1):45–60, Feb 2020.

6. A. Fernandez, W. Deng, S. L. McLaughlin, A. C. Pirkey, S. L. Rellick, A. Razazan, and D. J. Klinke. Cell Communication Network factor 4 promotes tumor-induced immunosuppression in melanoma. EMBO Rep, 23(4):e54127, Apr 2022.

7. D. J. Klinke and A. Torang. An unsupervised feature extraction and selection strategy for identifying epithelial-mesenchymal transition state metrics in breast cancer and melanoma. iScience, 23(5):101080, 2020.

8. D. J. Klinke, A. Fernandez, W. Deng, A. Razazan, H. Latifizadeh, and A. C. Pirkey. Data-driven learning how oncogenic gene expression locally alters heterocellular networks. Nat Commun, 13(1):1986, Apr 2022.

9. K. Gibby, W. K. You, K. Kadoya, H. Helgadottir, L. J. Young, L. G. Ellies, Y. Chang, R. D. Cardiff, and W. B. Stallcup. Early vascular deficits are correlated with delayed mammary tumorigenesis in the MMTV-PyMT transgenic mouse following genetic ablation of the NG2 proteoglycan. Breast Cancer Res., 14(2):R67, Apr 2012.

10. D. I. Gabrilovich, S. Ostrand-Rosenberg, and V. Bronte. Coordinated regulation of myeloid cells by tumours. Nat. Rev. Immunol., 12(4):253–268, March 2012.

11. V. Bronte and M. J. Pittet. The spleen in local and systemic regulation of immunity. Immunity, 39(5):806–818, November 2013.

12. M. H. Kubala, V. Punj, V. R. Placencio-Hickok, H. Fang, G. E. Fernandez, R. Sposto, and Y. A. DeClerck. Plasminogen activator inhibitor-1 promotes the recruitment and polarization of macrophages in cancer. Cell reports, 25(8):2177–2191.e7, 2018.

13. Wang D, Yang L, Yu W, Wu Q, Lian J, Li F, Liu S, Li A, He Z, Liu J, Sun Z, Yuan W, and Zhang Y. Colorectal cancer cell-derived ccl20 recruits regulatory t cells to promote chemoresistance via foxo1/cebpb/nf-?b signaling. J Immunother Cancer, 7(1):215, 2019.

14. J. Korbecki, K. Barczak, I. Gutowska, D. Chlubek, and I. Baranowska-Bosiacka. Cxcl1: Gene, promoter, regulation of expression, mrna stability, regulation of activity in the inter-cellular space. Int J Mol Sci, 23(2), 2022.

15. R. Reschke, J. Yu, B. Flood, E. F. Higgs, K. Hatogai, and T. F. Gajewski. Immune cell and tumor cell-derived cxcl10 is indicative of immunotherapy response in metastatic melanoma. J Immunother Cancer, 9(9):e003521, 2021.

16. D. J. Klinke, N. Horvath, V. Cuppett, Y. Wu, W. Deng, and R. Kanj. Interlocked positive and negative feedback network motifs regulate β-catenin activity in the adherens junction pathway. Mol. Biol. Cell, 26(22):4135–4148, Nov 2015.

17. G. Berx, A. M. Cleton-Jansen, K. Strumane, W. J. de Leeuw, F. Nollet, F. van Roy, and C. Cornelisse. E-cadherin is inactivated in a majority of invasive human lobular breast cancers by truncation mutations throughout its extracellular domain. Oncogene, 13(9):1919– 1925, Nov 1996.

18. M. Austinat, R. Dunsch, C. Wittekind, A. Tannapfel, R. Gebhardt, and F. Gaunitz. Correlation between beta-catenin mutations and expression of Wnt-signaling target genes in hepatocellular carcinoma. Mol Cancer, 7:21, Feb 2008.

19. M. Iwamoto, D. J. Ahnen, W. A. Franklin, and T. H. Maltzman. Expression of beta-catenin and full-length APC protein in normal and neoplastic colonic tissues. Carcinogenesis, 21 (11):1935–1940, Nov 2000.

20. A. Leask. Conjunction junction, what’s the function? CCN proteins as targets in fibrosis and cancers. Am. J. Physiol., Cell Physiol., 318(6):C1046–C1054, Jun 2020.

21. Y. M. Kulkarni, E. Chambers, A. J. R. McGray, J. S. Ware, J. L. Bramson, and D. J. Klinke. A quantitative systems approach to identify paracrine mechanisms that locally suppress immune response to interleukin-12 in the b16 melanoma model. Integr Biol (Camb), 4: 925–36, 2012.

22. D. J. Klinke. An age-structured model of dendritic cell trafficking in the lung. Am J Physiol Lung Cell Mol Physiol, 291:1038, 2006.

23. D. J. Klinke. A multi-scale model of dendritic cell education and trafficking in the lung: Implications for t cell polarization. Ann Biomed Eng, 35:937–955, 2007.

24. C. S. Garris, S. P. Arlauckas, R. H. Kohler, M. P. Trefny, S. Garren, C. Piot, C. Engblom, C. Pfirschke, M. Siwicki, J. Gungabeesoon, G. J. Freeman, S. E. Warren, S. Ong, E. Browning, C. G. Twitty, R. H. Pierce, M. H. Le, A. P. Algazi, A. I. Daud, S. I. Pai, A. Zippelius, R. Weissleder, and M. J. Pittet. Successful Anti-PD-1 Cancer Immunotherapy Requires T Cell-Dendritic Cell Crosstalk Involving the Cytokines IFN-γ and IL-12. Immunity, 49(6): 1148–1161, Dec 2018.

25. D. J. Klinke and K. M. Brundage. Scalable analysis of flow cytometry data using R/Bioconductor. Cytometry A, 75(8):699–706, Aug 2009.

26. S. M. Lin, P. Du, W. Huber, and W. A. Kibbe. Model-based variance-stabilizing transformation for illumina microarray data. Nucleic acids research, 36(2):e11, 2008.

